# Potential probiotic application of a novel commensal *E. coli* with antagonistic activity against different enteric pathogens

**DOI:** 10.64898/2025.12.03.692057

**Authors:** Prolay Halder, Sanjib Das, Soumalya Banerjee, Arindam Mukherjee, Supriya Mandal, Ashis Debnath, Nivedita Roy, Asish Kumar Mukhopadhyay, Debaki Ranjan Howlader, Hemanta Koley

## Abstract

Diarrheal diseases remain a leading cause of global morbidity and mortality, and treatment options are increasingly compromised by the global rise in multidrug-resistant (MDR) pathogens. Furthermore, the absence of effective vaccines against many causative agents renders large populations vulnerable. This critical gap necessitates the development of novel alternatives that can function both prophylactically to prevent infection and therapeutically to treat established diseases. Here, we investigated the commensal *Escherichia coli* strain HK220822 (HK5), which was isolated from a healthy human, as a potential live biotherapeutic. Comprehensive genomic and phenotypic characterization established a robust safety profile, confirming the absence of key virulence genes while demonstrating essential probiotic traits, including high tolerance to gastrointestinal stressors, such as acid and bile. The functional efficacy of *Escherichia coli* HK5 has been validated in murine models of infection. Prophylactic administration before pathogen exposure significantly inhibited intestinal colonization by *Salmonella* Typhimurium, *Shigella flexneri 2a*, and *Vibrio cholerae* (El Tor O1 strain). Moreover, HK5 demonstrated therapeutic potential by significantly reducing pathogen shedding when administered to previously infected animals. These findings establish commensal *E. coli* (HK5) as a potent candidate with dual preventative and therapeutic efficacy, offering a promising non-antibiotic strategy to combat diarrheal diseases.

## Introduction

Diarrheal diseases represent a persistent and formidable challenge to global public health, particularly in developing nations, where access to clean water and sanitation is limited. Annually, these infections account for over 1.7 billion episodes of illness and are a primary cause of mortality in children under five years of age, claiming approximately 525,000 young lives each year (1). This immense burden is quantified by Disability-Adjusted Life Years (DALYs), reflecting millions of years of healthy life lost to premature death and disability (2). In India, the situation is especially critical, with diarrheal diseases constituting a leading cause of pediatric hospitalization and mortality. A substantial portion of this burden is driven by highly virulent enteric bacterial pathogens including *Salmonella Typhimurium*, *Shigella flexneri* 2a, and *Vibrio cholerae*. The clinical management of these infections is now severely threatened by the escalating crisis of multidrug resistance (MDR), which renders established antibiotic regimens increasingly ineffective and increases the risk of treatment failure, prolonged illness, and death (3–6).

An ideal strategy for combating these pathogens is widespread and effective vaccination. However, this approach also has significant limitations. Despite decades of intensive research, no broadly licensed and effective vaccines are currently available to protect against diverse serovars of non-typhoidal *Salmonella* or *Shigella* (7–9). Although oral cholera vaccines (OCVs) for *Vibrio cholerae* exist, their public health impact remains constrained by considerable logistical and implementation challenges (10–11). Factors such as the required two-dose schedule, difficulties in maintaining a cold chain in resource-poor settings, low public awareness outside active outbreaks, and inconsistent vaccine supply chains frequently lead to poor uptake and incomplete community coverage, leaving large populations vulnerable. Although oral rehydration therapy (ORT) is a life-saving cornerstone for managing dehydration, it is a supportive measure. ORT replenishes fluids and electrolytes but does not address the underlying etiology of the infection, failing to eliminate the pathogen, shortening the duration of illness, or curbing pathogen shedding and transmission (12–13).

In this study, we characterized a novel commensal *Escherichia coli* strain, HK220822 (HK5), isolated from a healthy human volunteer. Comprehensive genomic and PCR-based screening confirmed the isolate to be non-pathogenic, lacking key virulence genes associated with diarrheagenic pathotypes, while demonstrating essential probiotic traits, such as high tolerance to acid and bile and robust biofilm formation. Functionally, *E. coli* HK220822 (HK5) exhibits selective *in vitro* antagonism against *Shigella flexneri*. Although a short-term co-infection model did not show significant pathogen reduction, the isolate demonstrated profound protective effects in more clinically relevant scenarios. In a murine model, prophylactic administration of *E. coli* HK220822 (HK5) one day prior to challenge significantly inhibited subsequent colonization by *Salmonella* Typhimurium, *Shigella flexneri*, and *Vibrio cholerae*. Moreover, *E. coli* HK220822 (HK5) acted therapeutically, and significantly reduced pathogen loads when administered to previously infected animals. These findings established HK5 as a potent probiotic candidate with both preventative and therapeutic efficacy against a range of clinically important enteric pathogens.

## Methods

### Ethical statement

All animal experiments were performed in compliance with (Government of India) guidelines and approved by the Institutional Animal Ethics Committee (IAEC) of the Indian Council of Medical Research – National Institute for Research in Bacterial Infections (ICMR-NIRBI, Registration No. 68/GO/ReBi/S/1999/CPCSEA valid 09/07/2029). Euthanasia was conducted according to the AVMA Guidelines for the Euthanasia of Animals (2020), and the study adheres to the ARRIVE guidelines. When required, the mice were euthanized with CO_2_. The machine was calibrated to operate at an inlet pressure of 345 kPa (50 PSIG) to ensure controlled and humane gas delivery; without pre-charging the chamber, the animals were placed inside, and 100% CO was introduced at a fill rate of 40% of the chamber volume per minute. For experiments requiring oral gavage, animals were sedated with ketamine (15 mg/kg body weight) mixed with xylazine (1 mg/kg body weight) and monitored during the experiment for signs of distress.

Stool samples were collected from healthy volunteers after informed written consent was obtained. This study was approved by the Institutional Ethical Committee (IEC) and Indian Council of Medical Research, India (human ethics approval no. A-1/2021-IEC, animal ethics approval no. PRO/184/ - Nov 2021-24).

### Animals

Female BALB/c mice (8–10 weeks old) were obtained from the animal facility of the ICMR-NIRBI, Kolkata. The mice were maintained in a specific-pathogen-free (SPF) facility under a regulated 12-hour light/dark schedule, with unrestricted access to sterile food and water. Animals were randomly allocated to the respective experimental groups. (n = 6 mice per group).

### Isolation and *in vivo* selection of a colonization-competent *Escherichia coli*

A human stool sample was obtained from a healthy volunteer and stored at –80°C. It was homogenized in sterile phosphate-buffered saline (PBS), subjected to serial dilutions, and subsequently spread onto MacConkey agar plates to select lactose-fermenting gram-negative bacteria. Following overnight incubation at 37°C, distinct lactose-positive (pink) colonies presumptive for *Escherichia coli* (*E. coli*), were purified by re-streaking on Luria-Bertani (LB) agar.

To select for robust gut colonization ability, an *in vivo* passage was performed. Briefly, a single pure colony was grown in Tryptic Soy Broth (TSB) to mid-logarithmic phase (OD₆₀₀ ≈ 0.6). Bacterial cells were harvested by centrifugation (4,000 × g, 10 min, 4°C), washed twice with sterile PBS, and resuspended in PBS to a final concentration of 1 × 10 CFU/mL. A 100 µL aliquot (1 × 10⁸ CFU) was administered to ten naïve BALB/c mice via oral gavage. All oral gavage experiments were preceded by administration of 5% NaHCO_3_ 15 min prior to bacterial challenge. After 24 h, fecal pellets were collected, suspended in PBS for homogenization, serially diluted, and cultured on LB agar plates. A dominant, well-isolated colony exhibiting morphology identical to that of the inoculum was selected, purified, and designated as strain HK220822 (a.k.a. HK5).

### Bacterial strains and culture conditions

The primary strains used in this study were the probiotic candidates *E. coli* HK220822 (HK5), *Shigella flexneri* 2a (2457T), *Salmonella enterica* ser. Typhimurium (S.17.7), and *Vibrio cholerae* O1 El Tor (N16961, El Tor, O1). For growth curve comparisons, *E. coli* Nissle 1917 (EcN) and *E. coli* ATCC 25922 were used as the reference strains. For PCR, Enterotoxigenic *E. coli* ETEC H10407, enterohaemorrhagic *E. coli* EHEC O157:H7, and Enteropathogenic *E. coli* (EPEC) 9327 were used. Enterohemorrhagic *E. coli* (EHEC) O157:H7 was also used for lipopolysaccharide (LPS) extraction. Unless otherwise specified, all strains were routinely cultured in TSB or on TSA at 37°C. Liquid cultures were grown under agitation (100 – 250 rpm), which was appropriate for the experiment. MacConkey, Hektoen Enteric (HE), and Thiosulfate-Citrate-Bile Salts-Sucrose (TCBS) agars were used to differentiate the specified strains based on their characteristic colony morphologies. Long-term stocks of all strains were maintained at –80°C in TSB containing 20% (v/v) glycerol.

### Whole-genome sequencing and bioinformatic analysis

WGS was performed on the Illumina platform to obtain a comprehensive genomic profile of the isolate, whole-genome sequencing (WGS) was performed on the Illumina platform. A genomic DNA library was prepared according to the manufacturer’s recommendations. The resulting raw sequence data were subjected to a rigorous bioinformatics analysis pipeline, as described below.

Initially, the quality of raw FASTQ reads was assessed using FastQC. The reads were then preprocessed for quality trimming and adapter removal using Fastp (v.0.23.4) (14). The processed high-quality reads were subsequently assembled *de novo* into contigs using Unicycler (v.0.4.4) (15). To improve the contiguity of the assembly, the contigs were scaffolded against a closely related reference genome using the RagTag (v2.1.0) (16). The overall quality, completeness, and statistical metrics of the final draft genome assembly were evaluated using QUAST (17).

For taxonomic confirmation at the species level, the 16S rRNA gene sequence was extracted from the final assembly using the ContEST16S tool within the EzBioCloud platform. The extracted sequence was compared against the EzBioCloud 16S database for identification (18, 19). The complete Whole Genome Shotgun project has been deposited in the GenBank database at the National Center for Biotechnology Information (NCBI) and is publicly available under accession number PQ305625.

### Phenotypic characterization of *E. coli* HK220822 (HK5)

To assess the phenotypic profile, *E. coli* HK220822 (HK5) was streaked onto MacConkey, HE, and Xylose Lysine Deoxycholate (XLD) agar, with LB agar serving as a non-selective growth control. Plates were incubated at 37°C for 24 h, and colony morphology and color changes were recorded to differentiate them from common enteric pathogens.

### Growth kinetics analysis

To compare the growth dynamics, *E. coli* HK220822 (HK5), *E. coli* Nissle 1917 (EcN), and *E. coli* ATCC 25922 were analyzed. Starter cultures were grown overnight in Tryptic Soy Broth at 37°C with shaking condition (200 rpm). These were then hundred times in flasks containing fresh, pre-warmed TSB. The Flasks were incubated at 37°C with continuous agitation (200 rpm). At hourly intervals for 12 hours, aliquots were aseptically removed for parallel measurement of optical density at 600 nm (OD₆₀₀) and viable cell counts. For viable counts, the samples were serially diluted in PBS and subsequently inoculated onto LB agar plates. Growth curves were generated by plotting OD₆₀₀ and log₁₀ CFU/mL against time.

### Assessment of acid and bile tolerance

The ability of this strain to survive harsh gastrointestinal conditions was evaluated by testing its tolerance to acidic pH and bile salts.

- **Acid tolerance:** An overnight culture of HK220822 (HK5) was inoculated (1% v/v) into 5 mL acidified TSB (pH 2.5) and a control TSB tube (pH 7.2).
- **Bile tolerance:** TSB was supplemented with 0.3% (w/v) ox bile salts (Sigma-Aldrich) at a physiologically relevant concentration and sterilized. An overnight culture of HK220822 (HK5) was inoculated (1% v/v) into a bile-supplemented TSB.

Both acid- and bile tolerance cultures were incubated for 3 h at 37°C with shaking (150 RPM). To quantify survival, liquid culture samples were collected at time zero and after the incubation period, serially diluted in PBS, and plated on LB agar. Viable cell counts (CFU/mL) were determined after overnight incubation at 37°C.

### Biochemical characterization of HK220822 (HK5)

HK220822 (HK5) was characterized using a standard panel of biochemical tests performed at 37°C. **Catalase activity** was assessed using 3% H₂O₂, where immediate effervescence indicated a positive result (20). **Urease production** was tested in Christensen’s Urea Broth, and a color change to cerise after 24 h was considered positive (21). For the **indole test**, Kovács’s reagent was added to a to 24-48-hour Tryptone Broth culture, with a superficial red ring confirming production (21). Carbohydrate utilization and H₂S production was evaluated on **Triple Sugar Iron (TSI) agar** slants inoculated by stabbing the butt and streaking the slant, with results recorded after 18–24 h (21). **Citrate utilization** was assessed by streaking a Simmons Citrate Agar slant, where a color change from green to Prussian blue after 18–48 h indicated a positive result (21). **Motility** was evaluated using the soft agar stab assay (LB with 0.3% agar). The isolate was stabbed into the center of the plate, with *Salmonella enterica* serovar Typhimurium and *Klebsiella pneumoniae* serving as the positive and negative controls, respectively. Diffuse turbidity radiating from the stab line after 16 h of incubation confirmed the motility.

### Antibiotic susceptibility testing

The antibiotic susceptibility profile of HK220822 (HK5) was determined using the Kirby-Bauer disk diffusion method following the guidelines of the Clinical and Laboratory Standards Institute (CLSI) (22). An inoculum was prepared by suspending the colonies in sterile saline to match the turbidity of the 0.5 McFarland standard. This suspension was used to create a confluent lawn on Mueller-Hinton agar (MHA) plates with a sterile cotton swab. The antibiotic disks were placed on the inoculated surfaces. plates were incubated at 37°C for 18–24 h under aerobic conditions. The diameters of the resulting zones of inhibition were measured, and the isolates were classified as susceptible (S), intermediate (I), or resistant (R), based on the breakpoints defined by the CLSI guidelines. The reference strain, *E. coli* ATCC 25922, was included in all assays for quality control. HK220822 (HK5) was tested against a panel of antibiotics representing multiple classes, including cephalosporin ceftriaxone (30 µg), carbapenem imipenem (10 µg), penicillin methicillin (30 µg), fluoroquinolones levofloxacin (5 µg), norfloxacin (10 µg), aminoglycosides neomycin (30 µg), and streptomycin (10 µg); quinolone nalidixic acid (30 µg); amphenicol chloramphenicol (30 µg); and folate synthesis inhibitors co-trimoxazole (25 µg) and trimethoprim (30 µg).

### Genotypic characterization by PCR-based screening

To confirm the identity of isolate HK5 and to establish its safety profile, genomic DNA (gDNA) was screened for the 16S rRNA gene and a panel of key virulence-associated genes. gDNA was isolated from overnight cultures using a standard phenol-chloroform extraction protocol.

PCR assays were performed to detect the conserved **16S rRNA gene** and screen for the absence of virulence genes common to enteric pathogens.

***ipaH*** (invasion), ***lt*** (enterotoxin), ***stx1*** and ***stx2*** (Shiga toxins), and ***eaeA*** (intimin). Primer sequences and PCR programs are listed in Supplemental Tables ST1 and ST2 (23, 24, 25, and 26). Reactions (25 µL) contained approximately 50 ng of gDNA, 0.5 µM of each primer, and Taq DNA polymerase in 1X reaction buffer with 200 µM dNTPs. Thermal cycling was carried out beginning with an initial denaturation at 95 °C for 5 min. This was followed by 30 amplification cycles, each comprising denaturation at 95 °C for 30 s, primer-dependent annealing for 30 s, and extension at 72 °C for 1 min. The reaction was completed in a final elongation step at 72 °C for 7 min. The amplified products were separated using agarose gel electrophoresis and a 2.0% (w/v) agarose gel pre-stained with ethidium bromide. and sized against an appropriate DNA ladder, namely, a one Kb GeneRuler DNA Ladder (cat. #. SM0311, Thermo Fisher Scientific) for STX-1 and -2, a 100 bp GeneRuler DNA Ladder (cat. # SM0241, Thermo) for LT, 16s rRNA, and Eae genes and a 100 bp Plus GeneRuler DNA Ladder (cat. # SM0321, Thermo Fisher Scientific) for LacY and IpaH.

A comprehensive set of controls was included in all the assays. *E. coli* ATCC 25922 and *E. coli* Nissle 1917 served as positive controls for 16S rRNA and negative controls for virulence genes. The following pathogenic strains were used as positive controls for specific virulence loci: *Shigella flexneri* 2a 2457T (*ipaH*), ETEC H10407 (*lt*), EHEC O157:H7 (*stx1* and *stx2*), and EPEC 9327 (*eaeA*). Nuclease-free water served as a no-template control.

### Isolation and analysis of lipopolysaccharide (LPS)

To characterize and compare the lipopolysaccharide (LPS) structure of HK220822 (HK5) against known pathogenic and commensal strains, LPS was extracted from HK220822 (HK5), EHEC O157:H7, *Shigella flexneri* 2a 2457t, and EcN. LPS was extracted from 500 mL of overnight culture using the hot phenol method (27). After cell lysis at 65°C, the aqueous phase was collected and LPS was precipitated with cold ethanol. The purified LPS pellet was resuspended in sterile water and quantified using the colorimetric carbohydrate assay (490 nm). Finally, equal amounts of LPS were resolved using 15% SDS-PAGE to distinguish between smooth (S-LPS) and rough (R-LPS) phenotypes. Following electrophoresis, the gel was subjected to a sensitive silver staining protocol involving periodic acid oxidation to visualize LPS bands (28). The stained gel was then imaged to compare the profiles and distinguish between smooth (S-LPS) and rough (R-LPS) phenotypes.

### Quantification of biofilm formation

The biofilm-forming capacity of HK220822 was quantified using a crystal violet microtiter plate assay with *EcN* and ATCC 25922 as controls. Overnight cultures were diluted 1:100 and incubated statically in 96-well plates at 30°C and 37°C for 24 and 72 h. After incubation, the wells were washed with PBS to remove the planktonic cells. Adherent biofilms were stained with 0.1% crystal violet and the bound dye was solubilized with 95% ethanol. Biofilm mass was quantified by measuring the absorbance of the solubilized stain at 595 nm.

### Assessment of antagonistic activity

To determine the ability of HK220822 (HK5) to inhibit the growth of enteric pathogens, its antagonistic properties were evaluated using in vitro and in vivo competition models.

- ***in vitro* co-culture assay:** The direct competitive activity of HK220822 (HK5) was tested against *S.* Typhimurium (S.17.7), *S. flexneri* 2a (2457T), and *V. cholerae* O1 El Tor (N16961). Overnight cultures of each strain were grown in TSB at 37°C. Equal volumes of an overnight culture of HK220822 (HK5) and one of the respective pathogens were combined in a fresh TSB. The co-cultures were incubated for 4 h at 37°C with agitation. Following incubation, the mixed bacterial population was harvested by centrifugation (4000 × g, 10 min, 4°C), serially diluted in sterile PBS, and plated onto selective media (n = 4 technical repeats for each of the three biological repeats) for differential enumeration. Viable counts for pathogens were determined on Hektoen Enteric (HE) agar (*S. flexneri*), Xylose Lysine Deoxycholate (XLD) agar (*S. Typhimurium*), and TCBS agar (*V. cholerae*), whereas HK220822 (HK5) was enumerated on MacConkey agar. Plates were incubated at 37°C and colony-forming units (CFU/mL) were counted.
- ***in vivo* co-infection model:** To evaluate competitive inhibition in the host environment, three groups of BALB/c mice (n = 6 per group) were used. Each group was challenged via oral gavage with a co-infection of HK220822 (HK5) with either *S.* Typhimurium (S.17.7), *S. flexneri* 2a (2457T), or *V. cholerae* (N16961). These co-cultures were prepared as described above where a four-hour old inoculum (1 × 10^8^ CFU/100 μl/mouse) was used in each case. After 24 h, fecal pellets were collected from each mouse to determine the extent of bacterial colonization. Fecal samples were weighed, homogenized in sterile PBS, serially diluted, and plated on appropriate selective media to differentially quantify each bacterial species. For example, fecal samples from mice colonized with HK5 and *Salmonella* typhimurium were plated on MacConkey and XLD agar plates. Colonies were compared and counted after incubation. Bacterial loads were calculated as CFU per gram of feces.

### Evaluation of Prophylactic and Therapeutic Efficacy of *E. coli* HK5

To assess the ability of *E. coli* HK220822 (HK5) to prevent and treat enteric pathogenic infections, in vivo prophylactic and therapeutic mouse models were established. For all in vivo challenges, the inoculum was prepared from mid-logarithmic-phase cultures. Briefly, single colonies of *E. coli* HK5 and the pathogenic strains were used to inoculate TSB and grown overnight at 37°C. The overnight cultures were then sub-cultured (1:100) into fresh TSB and grown to the mid-logarithmic phase. Bacterial cells were harvested by centrifugation (8,000 × g, 10 min, 4°C), washed twice with sterile PBS, and finally resuspended in PBS to the required concentrations for oral gavage.

- **Prophylactic Model:** To evaluate the protective potential of HK5, mice (n = 6 per group) were pre-colonized with the probiotic candidate prior to pathogen exposure. On day -1, mice received a single oral gavage of 1 × 10⁸ CFU of *E. coli* HK5 in 100 µL of PBS. The parallel control group was administered an equal volume of sterile PBS. On day 0, all mice were challenged via oral gavage with an infectious dose (1 × 10⁷ CFU/mouse) of either *S. Typhimurium* (S.17.7), *S. flexneri* 2a (2457T), or *V. cholerae* (N16961).
- **Therapeutic Model:** To assess the ability of HK5 to treat an established infection, mice (n = 6 per group) were first infected on day 0 via oral gavage with 1 × 10 CFU of *S. Typhimurium*, *S. flexneri*, or *V. cholerae*. Twenty-four hours later (day +1), the treatment group received a single therapeutic dose of 1 × 10⁸ CFU of *E. coli* HK5. The control group was administered sterile PBS at the same time points.

For both models, fecal pellets were collected 24 h after pathogen challenge (on day +1 for the prophylactic study and day +2 for the therapeutic study). Fecal samples were weighed, homogenized in sterile PBS, and serially diluted. To quantify bacterial populations differentially, dilutions were plated onto the appropriate selective media. Bacterial loads were calculated and reported as CFU per gram of feces.

### Statistical analyses

Data visualization and statistical analyses were performed using R (v4.2.1) with the tidyverse, ggpubr, and rstatix packages and GraphPad Prism (v. 8.0.2, GraphPad Software, Inc.). Quantitative data are presented as the mean ± standard error of the mean (SEM) or as boxplots depicting the median and interquartile range, as specified in the figure legends. For all *in vivo* experiments, a sample size of n six mice per group was used.

The choice of the statistical test was based on the experimental design and data distribution. For comparisons between two paired groups, such as *in vitro* co-culture and *in vivo* co-infection assays, the non-parametric Wilcoxon signed-rank test was used. For the multigroup biofilm assay, statistical analysis was performed using two-way ANOVA, and multiple comparisons were performed using Tukey’s test. Statistical significance was determined at a threshold of p < 0.05. The significance levels are denoted in the figures as follows: p < 0.05, *p < 0.01, and **p < 0.001.

## Results

### Genomic and Phylogenetic Analyses Identify Isolate HK220822 (HK5) as *Escherichia coli*

To determine the taxonomic identity of the isolate HK220822 (HK5), we first performed 16S rRNA gene sequencing. This initial approach yielded ambiguous results; a BLASTn search showed 100% sequence identity with numerous *Escherichia coli* strains, but also with several *Shigella* species. This reflects the well-established limitation of the 16S rRNA gene in differentiating between these closely related genera.

This ambiguity was underscored by phylogenetic analysis. A neighbor-joining tree generated using the NCBI BLAST algorithm placed HK5 within a broad, unresolved clade of *E. coli* strains (Fig. 1A). In contrast, a phylogenetic tree constructed using the EzBioCloud 16S database showed that HK5 clustered distinctly with *Shigella* species, positioned closest to *Shigella flexneri* (Fig. 1B).

**Figure 1.**
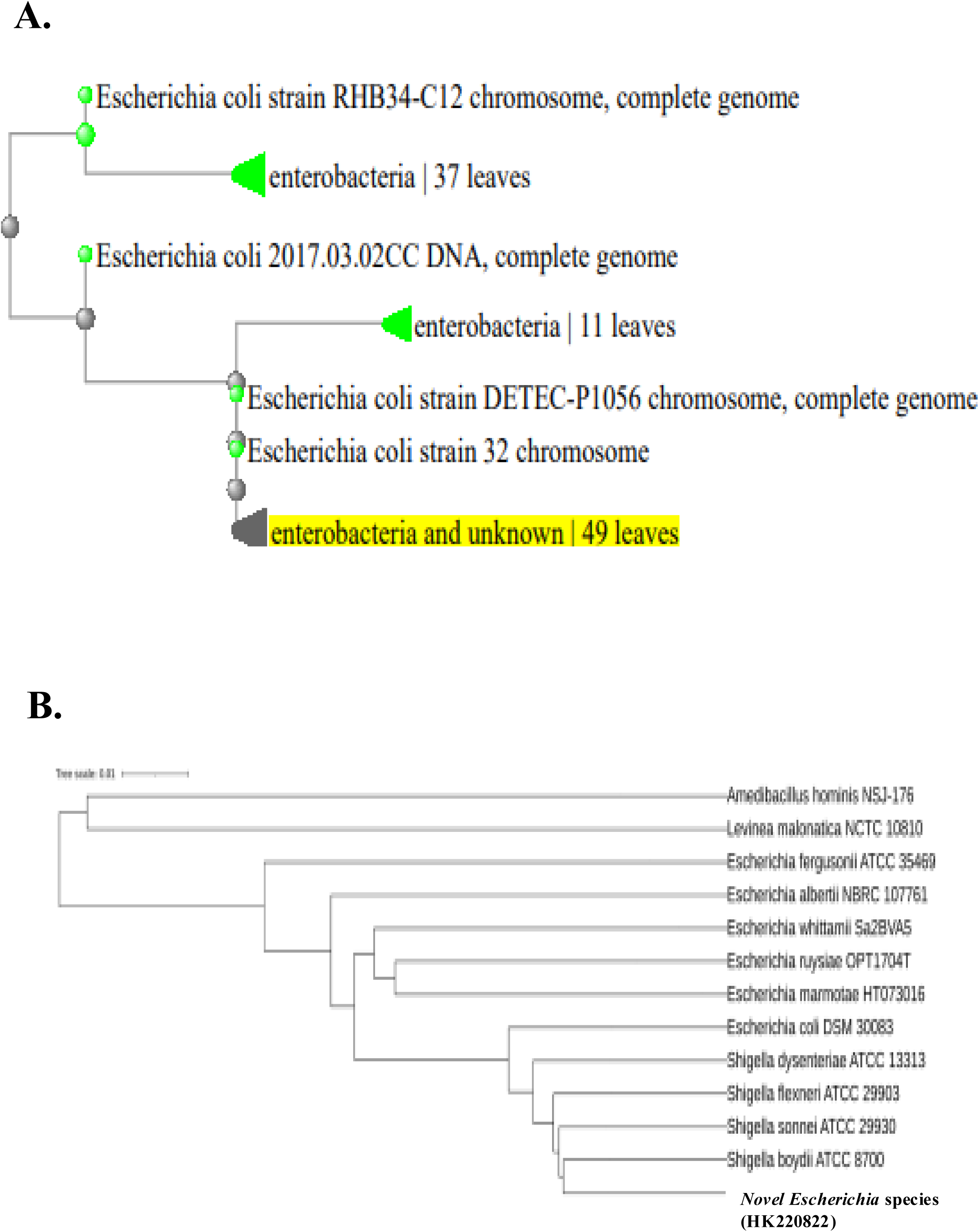
Phylogenetic analysis of *E. coli* HK220822 (HK5) based on the 16S rRNA gene sequence. **(A)** Phylogenetic tree generated from the NCBI BLASTn alignment. The position of the query sequence HK220822 (HK5) is indicated by the yellow highlight, showing its placement within a large clade dominated by *Enterobacteriaceae*, primarily *Escherichia coli* strains. **(B)** A formal phylogenetic tree constructed using the EzBioCloud platform, showing the relationship of HK220822 to its closest relatives. The isolate clusters within the *E. coli*/*Shigella* group, branching closely with species of the genus *Shigella* (*S. dysenteriae*, *S. flexneri*, *S. sonnei*, *S. boydii*) and *E. coli* DSM 30083. This clustering highlights the well-known limitation of the 16S rRNA gene in resolving the taxonomy of these two closely related genera.

To resolve this taxonomic uncertainty and obtain a definitive classification, we performed whole-genome sequencing (WGS) of the HK5. The *de novo* assembly produced a single circular chromosome of 5.17 Mb. Whole-genome analysis using Average Nucleotide Identity (ANI) definitively identified the isolate as *Escherichia coli*. HK5 shared 96.6% ANI with the *E. coli* type strain, a value comfortably above the 95% species-delineation threshold. This classification was further corroborated using the Genome Taxonomy Database Toolkit (GTDB-Tk).

A comprehensive overview of HK5 genomic architecture is presented in the circular genome map (Supplemental Fig. S1). This map illustrates the distribution of coding sequences (CDS) organized by functional clusters of orthologous groups (COG) categories, the location of tRNA and rRNA genes, and variations in GC content and GC skew across the chromosome.

### Phenotypic and Biochemical Profile of *E. coli* HK220822 (HK5)

Following genomic identification, we characterized the phenotypic and biochemical profiles of *E. coli* HK5. On standard Luria-Bertani (LB) agar, the strain formed smooth pale-yellow colonies (Fig. 2A). On selective and differential media, HK5 displayed characteristics consistent with those of an enteric bacterium capable of vigorous sugar fermentation. It produced vibrant pink colonies on MacConkey agar, confirming lactose fermentation (Fig. 2B), and bright orange-yellow colonies on Hektoen Enteric (HE) agar, indicating fermentation of sugars in the medium without H_2_S production (Fig. 2C). Similarly, it formed yellow colonies on Xylose Lysine Deoxycholate (XLD) agar owing to acid production from sugar fermentation (Fig. 2D).

**Figure 2.**
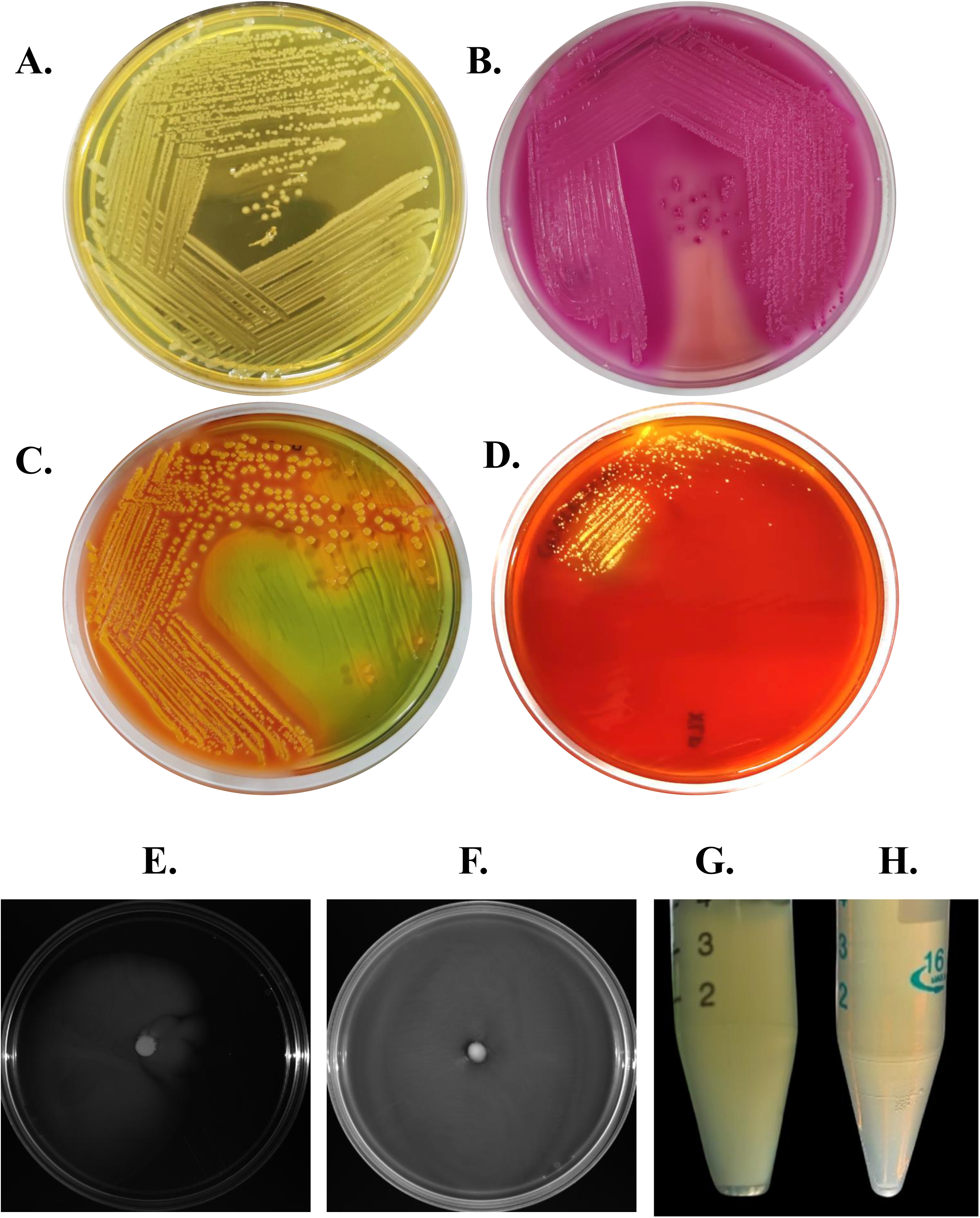
Phenotypic and physiological characterization of *E. coli* HK220822 (HK5). Colony morphology of HK5 on **(A)** Luria-Bertani (LB) agar, **(B)** MacConkey agar, showing characteristic pink colonies indicative of lactose fermentation, **(C)** Hektoen Enteric (HE) agar, showing orange-yellow colonies from sugar fermentation, and **(D)** Xylose Lysine Deoxycholate (XLD) agar, showing yellow colonies due to acid production. **(E, F)** Motility assay in soft agar demonstrating **(E)** the non-motile phenotype of HK5, with growth confined to the stab line, compared to **(F)** the diffuse turbidity of the motile control, *Salmonella* Typhimurium. **(G, H)** Survival and growth of HK5 after a 3-hour challenge in **(G)** acidic TSB (pH 2.5) and **(H)** TSB supplemented with 0.3% bile salts, indicated by the persistent turbidity of the cultures.

A panel of biochemical tests revealed an atypical metabolic profile of the *E. coli* strain. HK5 cells were catalase-positive but urease-negative. Notably, the tumor was indole-negative and citrate-positive. HK5 produced an acid slant and acid butt (A/A) on Triple Sugar Iron (TSI) agar, confirming its ability to ferment both glucose and lactose/sucrose, but did so without producing gas or H_2_S. In a soft agar stab assay, HK5 was found to be non-motile, with growth confined to the inoculation line (Fig. 2E), in stark contrast with the diffuse growth pattern of the motile control strain (Fig. 2F). Complete biochemical profiles are summarized in Supplemental Table ST3.

### HK220822 (HK5) Exhibits Robust Tolerance to Gastrointestinal Stress and Possesses a Favorable Antimicrobial Susceptibility Profile

A key attribute for a successful oral probiotic is the ability to survive transit through harsh conditions of the gastrointestinal tract. Therefore, we quantified the tolerance of *E. coli* HK5 cells to both acidic pH and bile salts. This strain demonstrated remarkable resilience, resulting in viability after a 3-hour exposure to a highly acidic environment (pH 2.5). Similarly, viability was retained following incubation in TSB supplemented with a physiologically relevant concentration of 0.3% ox bile. While these quantitative data confirm its robustness, visual evidence of dense culture growth following these stress challenges is provided in Figure 2G and 2H.

To assess the safety profile regarding antibiotic resistance, the antimicrobial susceptibility of HK220822 (HK5) was determined and compared to that of the reference strain *E. coli* ATCC 25922 and a representative commensal *E. coli* (Supplemental Table ST4). Although HK220822 (HK5) exhibited resistance to cefotaxime and methicillin and intermediate resistance to imipenem and neomycin, it remained susceptible to other key antibiotics, including trimethoprim and tetracycline. This profile suggests that HK220822 (HK5) does not harbor a broad multidrug-resistant phenotype, which is a critical safety consideration for probiotic candidates.

### Genotypic Profile of *E. coli* HK220822 (HK5) Confirms the Absence of Key Virulence Genes

To assess the safety profile of *E. coli* HK220822 (HK5) at the genetic level, PCR-based screening was conducted for species identity markers and a panel of major virulence genes associated with diarrheagenic *E. coli* and *Shigella* pathotypes.

First, the identities of the isolates were confirmed. *E. coli* HK220822 (HK5) tested positive for the *E. coli* 16S rRNA gene (Fig. 3A) and lactose permease gene (*lacY*), consistent with its lactose-fermenting phenotype (Fig. 3B). Subsequently, the strain was screened for virulence determinants and was found to be negative for all factors tested. It lacks the invasion plasmid antigen H gene (*ipaH*), a primary genetic marker for enteroinvasive *E. coli* (EIEC) and *Shigella* species (Fig. 3C). The strain was also negative for the heat-labile enterotoxin gene (*lt*), a hallmark of enterotoxigenic *E. coli* (ETEC) (Fig. 3D). Furthermore, *E. coli* HK220822 (HK5) did not possess the intimin gene (*eaeA*), which encodes a critical adhesion factor located on the locus of the enterocyte effacement (LEE) pathogenicity island found in enteropathogenic (*EPEC*) and enterohemorrhagic *E. coli* (EHEC) (Fig. 3E). Finally, the isolate tested negative for the genes responsible for encoding both Shiga toxin 1 (*stx1*) as well as Shiga toxin 2 (*stx2*), the definitive toxins of EHEC (Fig. 3F. i and 3F.ii, respectively).

**Figure 3.**
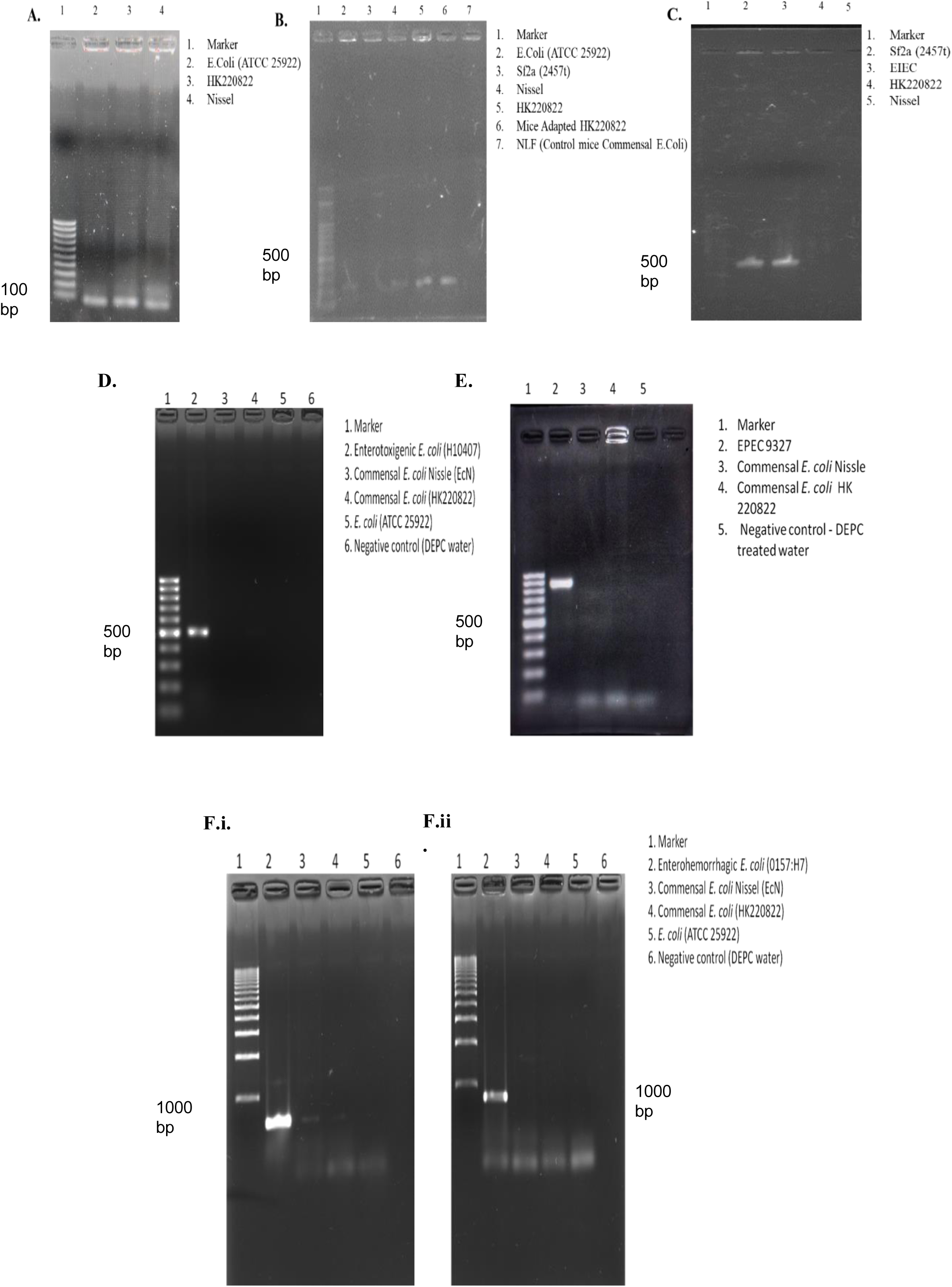
Genotypic screening confirms the identity of *E. coli* HK220822 (HK5) and demonstrates the absence of key virulence genes. Agarose gel electrophoresis of PCR amplicons. **(A)** Amplification of the 16S rRNA gene (∼86 bp) confirmed the identity of the isolate as *E. coli*. **(B)** Detection of the lactose-permease gene (*lacY*, ∼210 bp) confirmed its lactose-fermenting capability. Panels **(C-F)** screen for major virulence factors, for which HK5 tested negative in all cases. **(C)** Absence of the invasion plasmid antigen H gene (*ipaH*, ∼619 bp) with *S. flexneri* 2a (2457t) as the positive control. **(D)** Absence of the heat-labile enterotoxin gene (*lt*, ∼508 bp) with Enterotoxigenic *E. coli* (ETEC) H10407 as the positive control. **(E)** Absence of the intimin gene (*eaeA*, ∼881 bp) with Enteropathogenic *E. coli* (EPEC) 9327 as the positive control. **(F)** Absence of Shiga toxin genes **(F.i)** *stx1* (∼614 bp) and **(F.ii)** *stx2* (∼779 bp), with Enterohemorrhagic *E. coli* (EHEC) O157:H7 serving as the positive control. For all gels, a DNA ladder was loaded in the first lane, and *E. coli* Nissle 1917 and/or *E. coli* ATCC 25922 were used as non-pathogenic negative controls.

In all assays, the corresponding pathogenic strains served as positive controls and produced amplicons of the expected size, thus validating the experiments. Collectively, these genotypic data demonstrate that *E. coli* HK220822 (HK5) lacks the defining genetic determinants for the enteroinvasive, enterotoxigenic, enteropathogenic, and enterohemorrhagic pathotypes, supporting its classification as a nonpathogenic strain.

### *E. coli* HK220822 (HK5) Exhibits a Rough Lipopolysaccharide Phenotype Similar to Probiotic E. coli Nissle 1917

The structure of lipopolysaccharide (LPS), a major component of the gram-negative outer membrane, was analyzed to compare the isolate *E. coli* HK220822 with the pathogenic and commensal strains. Following extraction, LPS was separated by sodium dodecyl sulfate-polyacrylamide gel electrophoresis and detected by silver staining. The pathogenic strains *Enterohemorrhagic E. coli* O157:H7 and *Shigella flexneri* 2a displayed a high-molecular-weight, ladder-like banding pattern, a profile characteristic of smooth LPS (S-LPS) containing O-antigen side chains (Fig. 4A, lanes 2 and 5).

**Figure 4.**
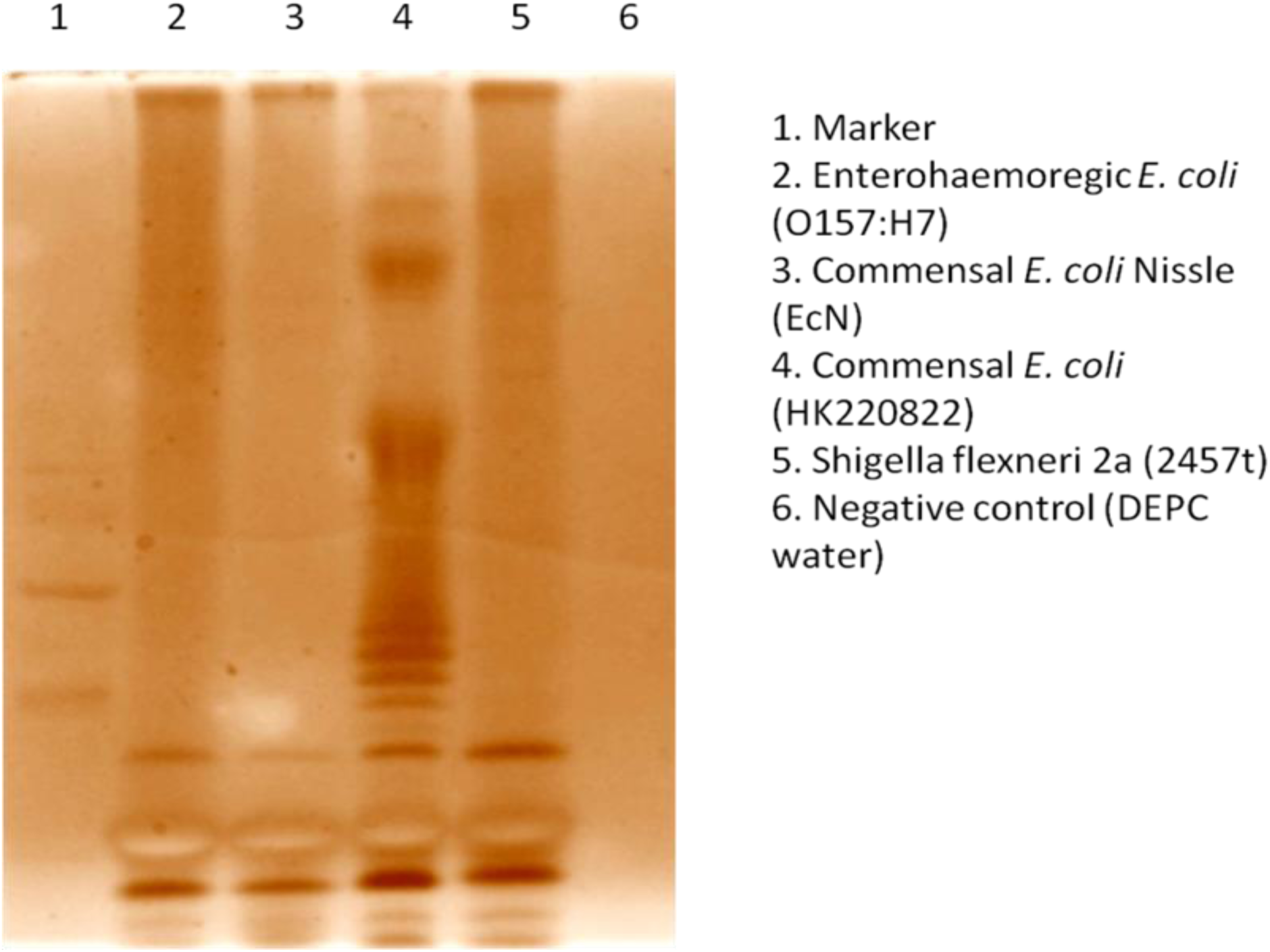
Analysis of the lipopolysaccharide (LPS) profile of *E. coli* HK220822 (HK5). Silver-stained SDS-PAGE gel of purified LPS. Lane 1 contains a molecular weight marker. LPS was loaded from *E. coli* O157:H7 (Lane 2), *E. coli* Nissle 1917 (EcN) (Lane 3), *E. coli* HK220822 (HK5) (Lane 4), and *Shigella flexneri* 2a (Lane 5). Lane 6 is a negative control.

In contrast, both *E. coli* HK220822 and the probiotic reference strain *E. coli* Nissle 1917 lacked this O-antigen ladder (Fig. 4A, Lanes 4 and 3, respectively). Their profiles were restricted to low-molecular-weight bands corresponding to the lipid A-core oligosaccharide. This is a defining feature of a rough LPS (R-LPS) phenotype. These findings demonstrate that *E. coli* HK220822 possesses rough-type LPS, a key surface characteristic that is shared with the probiotic *E. coli* Nissle 1917, distinguishing it from the S-LPS-producing pathogenic strains tested.

### Growth and Functional Characteristics of *E. coli* HK220822 (HK5)

To functionally characterize *E. coli* HK220822 (HK5) as a potential probiotic, its fundamental physiological properties, including growth dynamics, biofilm formation, and antimicrobial susceptibility, were assessed. These characteristics were benchmarked against the well-established probiotic *E. coli* Nissle 1917 and laboratory reference strain *E. coli* ATCC 25922.

The in vitro growth kinetics of the three strains were compared in a nutrient-rich broth for 24 h. *E. coli* HK220822 (HK5) exhibited growth dynamics that were comparable to those *E. coli* Nissle 1917 and ATCC 25922 (Fig. 5A). All strains demonstrated a similar lag phase, entered exponential growth between 2 and 3 h, and reached a stationary phase with a comparable cell density (OD < 1.0) at approximately 5 h post-inoculation. These data indicated that *E. coli* HK220822 (HK5) possesses robust growth characteristics typical of commensal *E. coli*.

**Figure 5.**
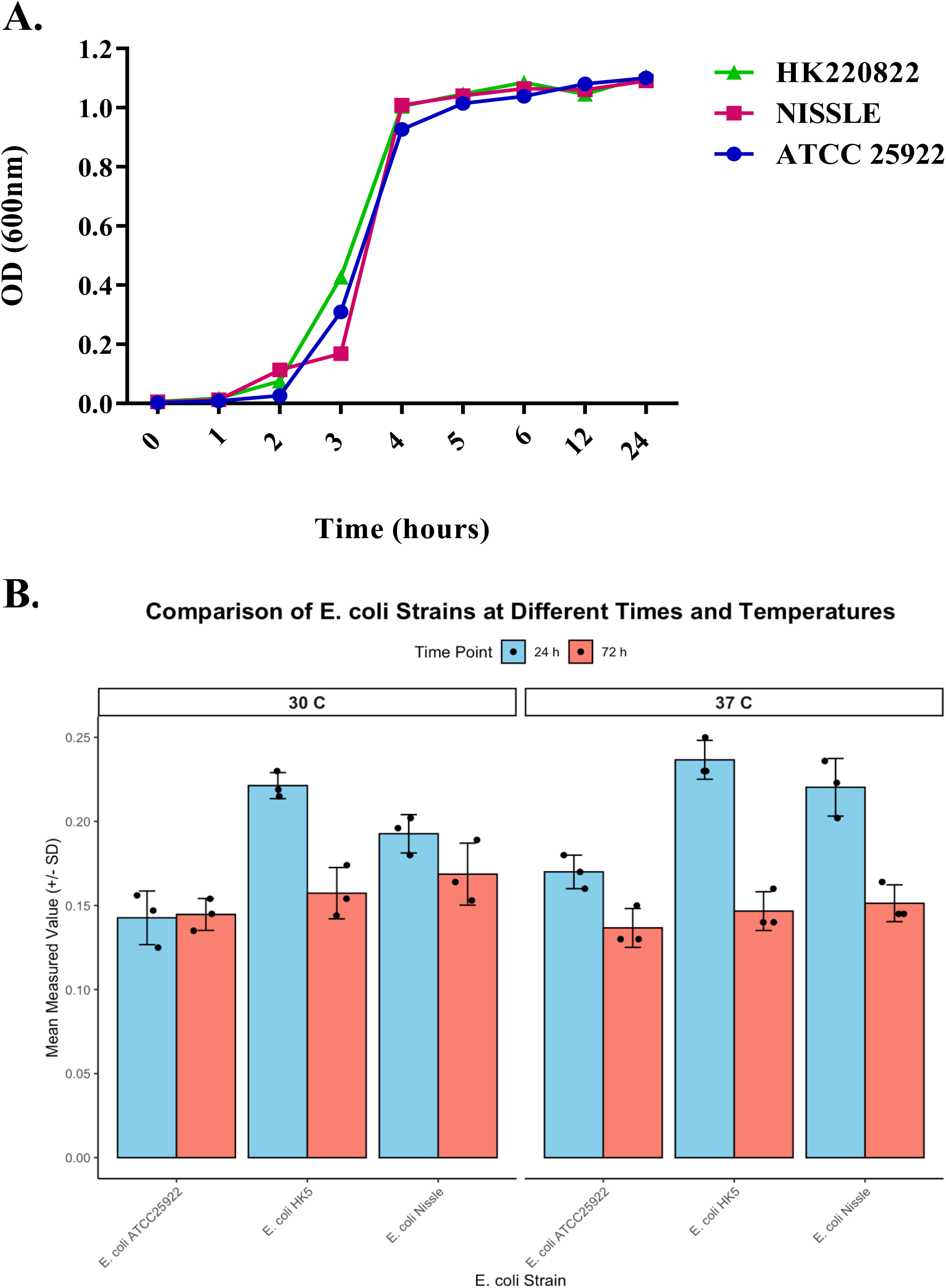
Growth kinetics and biofilm formation of *E. coli* HK220822 (HK5) compared to control strains. **(A)** Growth curves of *E. coli* HK220822 (HK5), *E. coli* Nissle 1917 (EcN), and *E. coli* ATCC 25922. Growth was monitored by measuring optical density at 600 nm (OD₆₀₀) over 24 hours in TSB at 37°C. Data points represent the mean of three independent experiments. **(B)** Quantification of biofilm formation by the three strains after 24 and 72 hours of static incubation at 30°C and 37°C. Biofilm biomass was quantified by crystal violet staining and measuring absorbance at 595 nm (shown on X-axis). Bars represent the mean of three replicates, and error bars show the standard deviation (SD). Individual data points from each replicate are overlaid as black dots. Statistical significance was determined using a Two-Way ANOVA followed by Tukey’s post-hoc test (see Supplemental Tables ST5 and ST6).

The ability to form biofilms, a key trait of intestinal colonization, was quantified at both 30°C and 37°C (Fig. 5B). At the 24-hour time point, both *E. coli* HK220822 (HK5) and *E. coli* Nissle demonstrated significantly greater biofilm formation than the laboratory strain, ATCC 25922, at both temperatures. Specifically, at 30°C, both HK5 (p < 0.001) and Nissle (p = 0.0019) produced more biofilms than ATCC 25922 (Supplemental Table ST5). A similar pattern was observed at 37°C, where both HK5 (p = 0.0001) and Nissle (p = 0.0008) again formed significantly more biofilms than ATCC 25922 (Supplemental Table ST6). After 72 h of incubation, no significant differences in biofilm mass were observed between the three strains at either temperature.

To further establish the safety profile, the antimicrobial susceptibility of *E. coli* HK220822 was determined. According to the antibiogram (Supplemental Table ST4), the isolate was resistant (R) to cefotaxime and methicillin and exhibited intermediate resistance (I) to imipenem and neomycin. Importantly, it remained susceptible (S) to other clinically relevant antibiotics, including trimethoprim and tetracycline.

### *E. coli* HK220822 (HK5) demonstrates selective in vitro antagonism against *Shigella flexneri* 2a (2457T)

To assess the direct competitive activity of *E. coli* HK5 cells, an in vitro coculture assay was performed. HK5 cells were co-incubated for 4 h with one of the three pathogenic strains: *Shigella flexneri* 2a (2457T) and *Salmonella enterica* ser. Typhimurium (S.17.7), or *Vibrio cholerae* O1 El Tor (N16961). Viable cell counts (CFU/mL) for both HK5 cells and the respective pathogens were determined using differential selective media.

The results showed that HK5 exerted a potent and specific inhibitory effect on the growth of *S. flexneri* 2a (2457T) (Figure 6A). Following 4-hour co-incubation, the viable count of *S. flexneri* was significantly lower than that of HK5 in the same cultures (p < 0.05). While HK5 maintained a high cell density (median CFU/mL > 10 < mL >), the *S. flexneri* population was suppressed by several orders of magnitude, with some technical replicates falling to the limit of detection.

**Figure 6.**
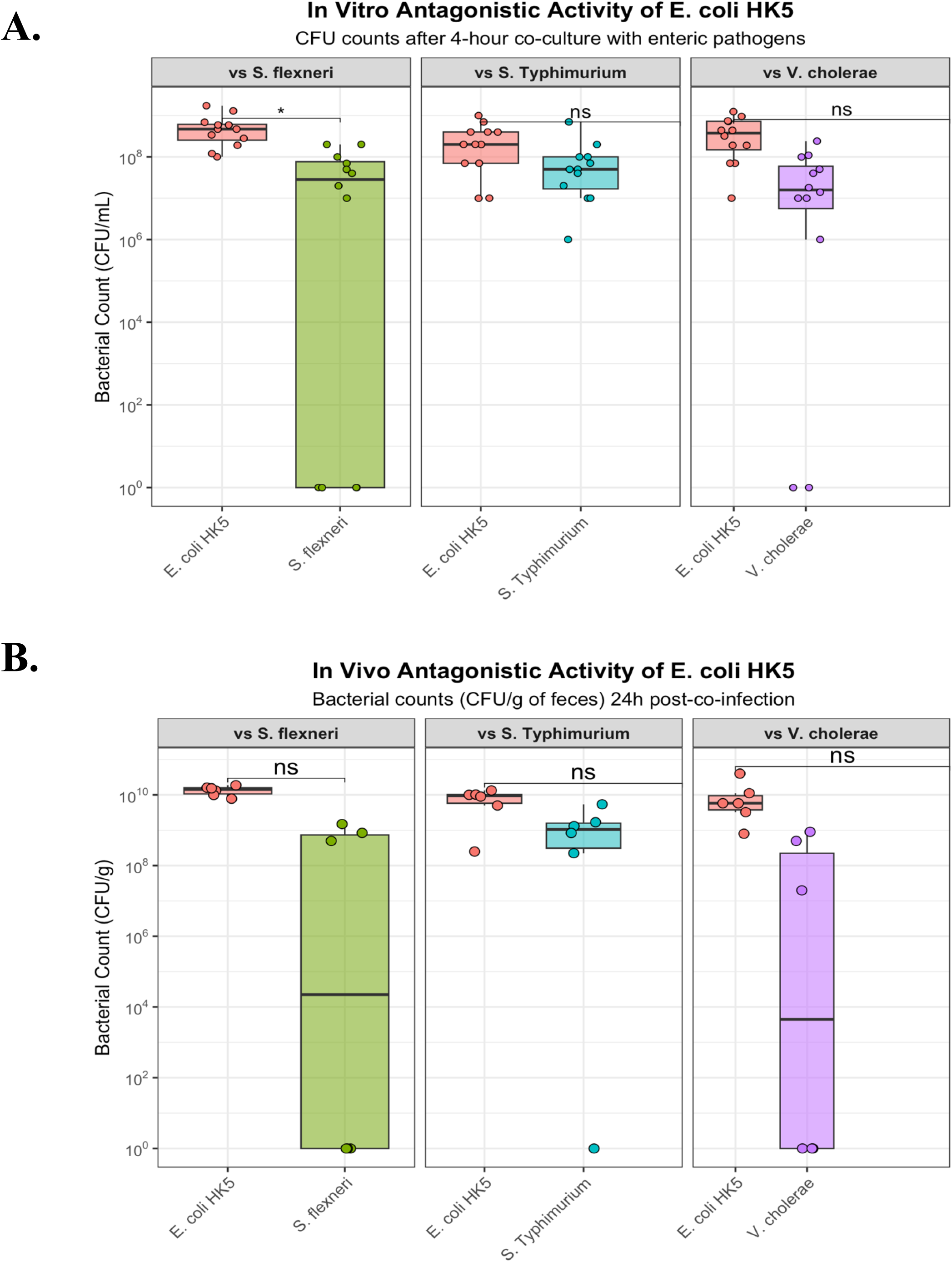
Antagonistic activity of *E. coli* HK220822 (HK5) (**A**) ***in vitro* co-culture assay reveals the antagonistic activity of *E. coli* HK5.** Bacterial counts (CFU/mL) were quantified after a 4-hour co-incubation of *E. coli* HK5 with the indicated pathogens. Each point represents a single measurement from a total of three biological experiments, each with four technical replicates. Boxplots show the median and interquartile range of the data. A paired t-test on log-transformed data was used for statistical comparison between HK5 and pathogen counts within each condition. **(B) *in vivo* co-infection does not reveal significant antagonism by *E. coli* HK5 against enteric pathogens.** BALB/c mice (n=6 per group) were co-infected via oral gavage with *E. coli* HK5 and one of three pathogens: *S. flexneri*, *S.* Typhimurium, or *V. cholerae*. Fecal bacterial loads were quantified 24 hours post-infection by selective plating and are expressed as Colony Forming Units per gram (CFU/g) on a logarithmic scale. The data are presented as boxplots where the central line indicates the median, the box edges represent the interquartile range, and all individual data points for each mouse are overlaid. A paired t-test was performed on log10-transformed data to compare the colonization levels of HK5 and the co-infected pathogen within each group (*p < 0.05; ns, not significant). ST: *Salmonella* Typhimuirum, Sf: *Shigella flexneri*, Vc: *Vibrio cholerae*.

In contrast, this antagonistic activity was not observed against any of the other tested pathogens. When co-cultured with *S.* Typhimurium (S.17.7) or *V. cholerae* (N16961), *E. coli* HK5 did not significantly reduce their viable counts (Figure 6A). In both conditions, HK5 and the pathogens grew to comparably high densities, and no statistically significant difference was found between their populations (p > 0.05).

These findings demonstrate that under the tested in vitro conditions, *E. coli* HK5 exhibited a targeted antagonistic interaction that significantly inhibited the growth of *S. flexneri* 2a but not *S.* Typhimurium or *V. cholerae*.

### *E. coli* HK220822 (HK5) colonizes robustly in vivo co-infection model

To determine whether the in vitro antagonism observed against *S. flexneri* would translate into a host environment, an in vivo co-infection model was employed. Groups of BALB/c mice (n = 6) were orally gavaged with a mixed inoculum of *E. coli* HK5 and one of three pathogens: *S. flexneri* 2a (2457T), *S.* Typhimurium (S.17.7), or *V. cholerae*(N16961). Fecal bacterial loads were quantified 24 h later to assess competitive colonization.

As shown in figure 6B, *E. coli* HK5 demonstrated robust colonization in the murine gut across all experimental groups, consistently reaching high densities of approximately 10–10 CFU/g of feces. The co-administered pathogens were also successfully colonized, although their levels were generally more variable than those of HK5. Notably, in several mice co-infected with *S. flexneri* or *V. cholerae*, the pathogen was below the limit of detection in the fecal samples at the 24-hour time point.

Despite its high level of colonization, paired statistical analysis found that *E. coli* HK220822 (HK5) did not significantly reduce the fecal counts of any of the tested pathogens. There was no statistically significant difference between the CFU/g of *E. coli* HK220822 (HK5) and that of the corresponding pathogen within any of the co-infection groups (*p* > 0.05). These results suggest that within the initial 24 h of this acute co-infection model, *E. coli* HK220822 (HK5) is an effective colonizer but does not exert a significant competitive exclusion or inhibitory effect on these pathogens in the gut.

### *E. coli* HK220822 (HK5) provides potent prophylactic and therapeutic protection in murine infection models

To determine the *in vivo* efficacy of *E. coli* HK220822 (HK5) against enteric pathogens, its ability to both prevent (prophylactic model) and treat (therapeutic model) infections was evaluated in mice.

In the prophylactic model, pre-colonization with a single dose of *E. coli* HK220822 (HK5) one day prior to infection conferred significant protection against all three pathogens tested (Figure 7, top panels). Twenty-four hours after challenge, the fecal loads of *S. flexneri* 2a, *S.* Typhimurium, and *V. cholerae* were all significantly lower than those of *E. coli* HK220822 (HK5) (*p* < 0.05 for all comparisons). This demonstrates that the prior establishment of *E. coli* HK220822 (HK5) in the gut effectively inhibits subsequent colonization by these pathogens.

**Figure 7:**
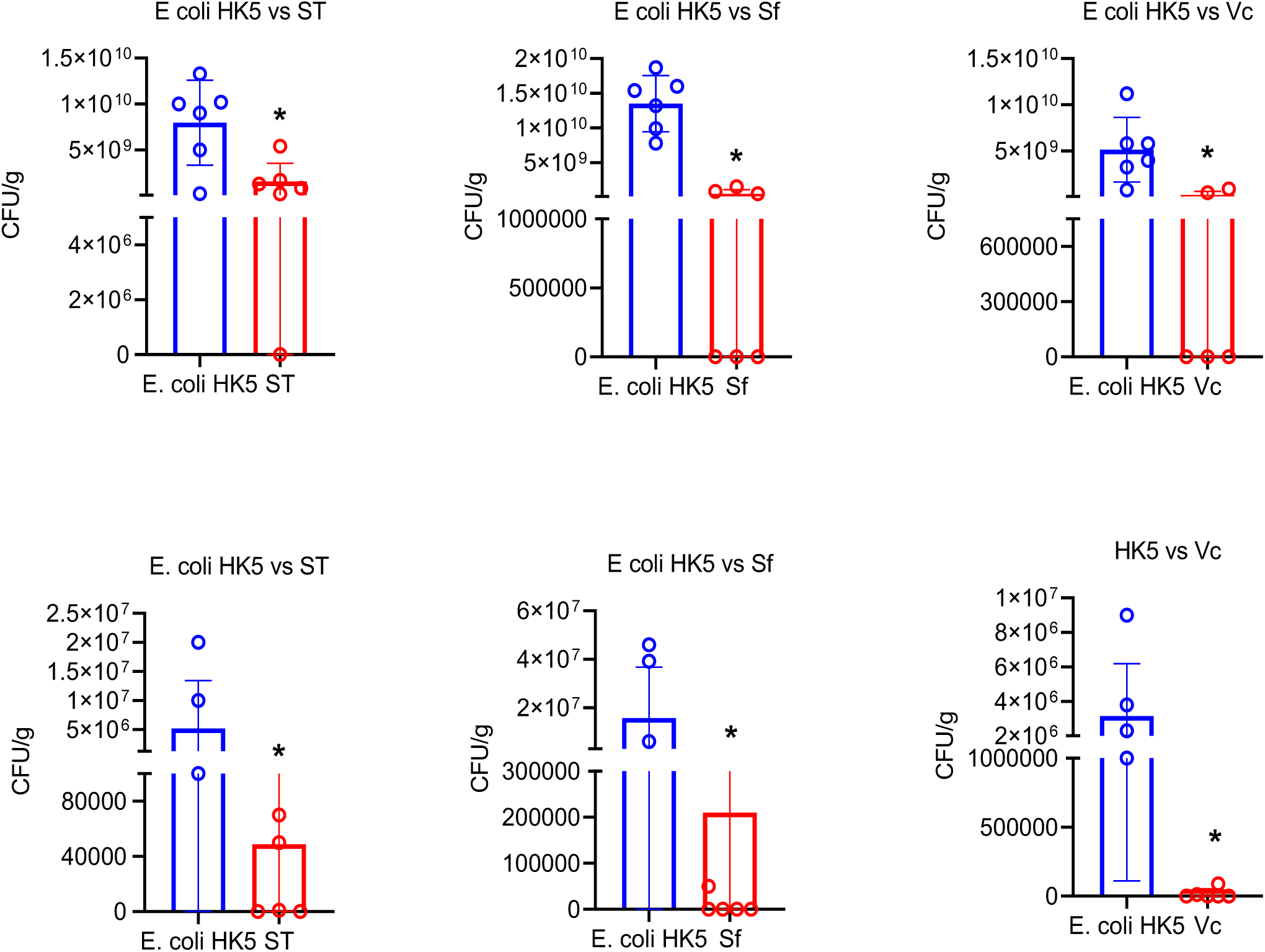
*E. coli* HK5 demonstrates significant prophylactic and therapeutic efficacy against enteric pathogens in vivo. The top panels show the prophylactic model, where mice were pre-treated with *E. coli* HK220822 (HK5) one day before challenge with *S. flexneri*, *S.* Typhimurium, or *V. cholerae*. The bottom panels show the therapeutic model, where mice were challenged with pathogens and treated 24 hours later with *E. coli* HK220822 (HK5). Fecal pathogen loads (CFU/g) were quantified 24 hours after challenge (prophylactic) or treatment (therapeutic). Data are presented as the mean ± standard deviation (SD), with points representing individual mice (n = 6 per group). Statistical significance between the control and HK5-treated groups was determined using a signed-rank test. All comparisons showed a significant reduction in pathogen load (p < 0.05).

Similarly, *E. coli* HK220822 (HK5) showed strong therapeutic activity against established infections (Figure 7, bottom panels). When administered 24 h after the mice had been infected with the pathogens, a single dose of *E. coli* HK220822 (HK5) significantly reduced fecal shedding of *S. flexneri*, *S.* Typhimurium, and *V. cholerae* compared to the control group that received PBS (*p* < 0.05).

Collectively, these findings show that *E. coli* HK220822 (HK5) is highly effective in the host environment, possessing both potent prophylactic capabilities to prevent infection and robust therapeutic activity to clear existing pathogens from the gut.

## Discussion

Diarrheal diseases continue to impose an immense global health burden, contributing significantly to morbidity and mortality, and staggering healthcare costs, particularly as measured by Disability-Adjusted Life Years (DALYs) (1). The escalating crisis of antimicrobial resistance and limitations of existing vaccines for many enteric pathogens have rendered conventional treatments increasingly inadequate. This critical gap underscores the urgent need for innovative non-antibiotic strategies.

Probiotic-based interventions, which leverage beneficial microbes to outcompete pathogens, represent a highly promising approach to reduce reliance on antibiotics and provide a sustainable solution for managing these devastating infections. The foundational trait of a successful oral probiotic is its ability to survive during transit through the gastrointestinal tract and establish itself within the complex gut ecosystem (29, 30, 31). Our initial characterization confirmed that the novel isolate *E. coli* HK220822 (HK5) possesses these essential characteristics. Its phenotypic profile on different media was consistent with that of a vigorous lactose-fermenting enterobacterium. Crucially, it demonstrated remarkable tolerance to both highly acidic conditions and physiologically relevant bile salt concentrations. This resilience is a non-negotiable prerequisite for any oral probiotic as it suggests that the bacterium can successfully navigate the harsh barriers of the stomach and small intestine to reach its site of action (30, 31).

Beyond mere survival, the safety and identity of probiotics must be unequivocally established. Along with the initial 16S rRNA sequencing, a definitive classification of HK220822 (HK5) as *E. coli* was achieved through whole-genome sequencing. This genomic-level analysis provides a framework for robust safety assessment. The absence of a suite of key virulence genes confirmed that HK5 lacks genetic determinants of major diarrheagenic pathotypes. This genotypic profile provides strong evidence that the strain is nonpathogenic and safe for potential host administration.

As expected, analysis of its surface lipopolysaccharide (LPS) revealed a rough phenotype, similar to the well-studied probiotic *E. coli* Nissle 1917, which has a semi-rough LPS thought to be key to its immunomodulatory properties. The ability of a probiotic to adhere to the mucosa and persist is fundamental for its function. The strong biofilm-forming capacity of *E. coli* HK220822 (HK5), which was significantly greater than that of the laboratory strain at 24 h, was a highly advantageous trait. Biofilm formation is a key mechanism for stable gut colonization, allowing the probiotic to create a protective niche, prolong its residence time, and compete more effectively with incoming pathogens for resources and attachment sites (32).

The functional efficacy of *E. coli* HK220822 (HK5) is multifaceted. While *in vitro* assays demonstrated potent and highly specific antagonism against *Shigella flexneri*, this direct inhibition was not replicated in a short-term *in vivo* co-infection model. This apparent discrepancy is highly informative, suggesting that the protective mechanisms of *E. coli* HK220822 (HK5) in the complex gut milieu are not based on simple direct killing. Instead, it points toward more sophisticated, time-dependent interactions, such as niche competition or modulation of the host environment (33). This hypothesis is strongly supported by the outcomes of prophylactic and therapeutic challenge models. When given a 24-hour head start to establish, *E. coli* HK220822 (HK5) provided profound prophylactic protection, significantly inhibiting the colonization of all three tested pathogens. Furthermore, when used to treat an established infection, *E. coli* HK220822 (HK5) acts therapeutically, leading to a significant reduction in pathogen shedding. The success of these clinically relevant models, in contrast to the neutral outcome of the acute co-infection experiment, strongly indicates that *E. coli* HK220822 (HK5) requires a period of engraftment to effectively condition the gut environment and exert its protective effects (34, 35).

In conclusion, this study identified and comprehensively validated *E. coli* HK220822 (HK5) as a promising probiotic candidate. It possesses a robust safety profile, essential physiological traits for gut survival and colonization, and, most importantly, demonstrated potent prophylactic and therapeutic efficacy against a range of clinically significant enteric pathogens. Its unique combination of features makes it a compelling subject for future mechanistic research. The proven functionality of *E. coli* HK220822 (HK5) in preclinical models establishes it as a strong candidate for the development of a novel live biotherapeutic to prevent and treat diarrheal diseases.

## Conclusion

This study identified commensal *Escherichia coli* HK220822 (HK5) as a promising live biotherapeutic agent for diarrheal diseases. Genomic and phenotypic analyses confirmed a robust safety profile, with HK5 lacking key virulence factors and demonstrating essential probiotic traits, such as high gastrointestinal stress tolerance and strong biofilm formation for effective gut colonization.

HK5 exhibited potent prophylactic and therapeutic efficacy in murine models against *Salmonella* Typhimurium, *Shigella flexneri*, and *Vibrio cholerae*. These protective effects, particularly pronounced after a period of engraftment, suggest mechanisms beyond direct antagonism that likely involve competitive exclusion and gut conditioning. The rough LPS phenotype further aligns with established immunomodulatory probiotics.

Given its validated safety, physiological fitness, and strong in vivo efficacy, HK5 is a compelling candidate for clinical development. Future studies should explore the mechanisms of action and translate these findings into human trials. This commensal *E*. *coli* HK5 strain offers a non-antibiotic strategy to address the urgent global health challenges associated with diarrheal illnesses.

## Acknowledgements

Prolay Halder gratefully acknowledges the Indian Council of Medical Research [ICMR-3/1/3/JRF-2018/HRD-066(66125)] for providing fellowship support during this study. Soumalya Banerjee [Student ID: 191620007740], Sanjib Das [SANJIB DAS/3336/(CSIR-UGC NET JUNE 2018)], and Arindam Mukherjee [Student ID: 231620010720] acknowledge the University Grants Commission for their fellowships, while the remaining authors express gratitude to the Indian Council of Medical Research for financial support. The authors also extend their thanks to Mr. Subrata Singha for his support in animal care and to Mr. Suhasit Ranjan Ghosh, Ms. Sristi Deb, Mr. Kaustav Adhikary, Ms. Usha Hansdah, and Mr. Rakesh Dey for their help with the technical work.

## Funding

This study was funded by the Indian Council of Medical Research under an institutional extramural project (Project Index. 5/9/1327/2020-Nut., RFC NO. NUT/ADHOC/19/2020-21). Dr. Debaki Ranjan Howlader is funded through DBT, India.

### Authors contributions

PH: Conceptualization, design, methodology, investigation, analysis, and writing; SD, SB, AM, SM, AD, NR: methodology and analysis; AKM: Review and editing; DRH: Data preparation, analysis, writing, and editing; HK: conceptualization, design, funding, supervision, writing, and editing. All authors have read and approved the final manuscript.

